# Kinetics of the inflammatory response during experimental *Babesia rossi* infection of beagle dogs

**DOI:** 10.1101/2021.09.16.460686

**Authors:** B.K. Atkinson, P. Thompson, E. Van Zyl, A. Goddard, Y. Rautenbach, J.P. Schoeman, V. Mukorera, A. Leisewitz

**Affiliations:** Department of Companion Animal Clinical Studies, Faculty of Veterinary Science, University of Pretoria, Pretoria, South Africa; Department of Production Animal Studies, Faculty of Veterinary Science, University of Pretoria, Pretoria, South Africa

## Abstract

*Babesia rossi* causes severe morbidity and mortality in dogs in sub-Saharan Africa, and the complications associated with this disease are likely caused by an unfocused, excessive inflammatory response. During this experimental *B. rossi* study we investigated inflammatory marker and cytokine kinetics during infection and after treatment. We aimed to determine whether infectious dose and treatment would influence the progression of the inflammatory response and clinical disease. Five healthy male beagle dogs were infected with *B. rossi*, three with a high infectious dose (HD group) and two with a low infectious dose (LD group). Clinical examination, complete blood count (CBC) and C-reactive protein (CRP) were determined daily. Cytokines were quantified on stored plasma collected during the study, using a canine specific cytokine magnetic bead panel (Milliplex©). The experiment was terminated when predetermined endpoints were reached. Parasitemia occurred on day 1 and 3 in the HD group and LD group respectively. The rate of increase in parasitemia in the HD group was significantly faster than that seen in the LD group. Significant differences were found in heart rate, blood pressure, interferon gamma (INFγ), keratinocyte chemoattractant (KC), INFγ-induced protein 10 (IP10), granulocyte-macrophage colony-stimulating factor (GM-CSF), monocyte chemoattractant protein 1 (MCP1), tumor necrosis factor alpha (TNFα), interleukin 2 (IL-2), IL-6, IL-7, IL-8, IL-10 IL-15, IL-18, CRP, neutrophils and monocytes between groups at multiple time points during the course of the infection. Our findings suggest that the initiation of inflammation occurs before the onset of clinical disease in *B. rossi* infection and infectious dose influences the onset of the inflammatory response. Treatment not only fails to curb the inflammatory response but may enhance it. Finally, we found that there is an imbalance in pro/anti-inflammatory cytokine concentrations during infection which may promote parasite replication.

## Introduction

*Babesia rossi*, a virulent *Babesia* species, causes a severe form of babesiosis in the domestic dog associated with a high rate of morbidity and mortality (1–3). Babesiosis is a complex multi-systemic disease that can be classified as either uncomplicated or complicated (2, 4, 5). Complicated babesiosis occurs when the pathology noted cannot be attributed purely to the anemia or when the anemia becomes severe enough to perpetuate organ dysfunction (2). *B. rossi* infection, like *Plasmodium falciparum* malaria in humans, results in a protozoal sepsis with a severe systemic inflammatory response (6–8). The concept of a ‘cytokine storm’ is well established in human inflammatory and infectious conditions, such as malaria and sepsis (9). This theory proposes that systemic illness and the course of disease is not solely caused by the microbes themselves but is also the result an unbalanced cytokine response to microbe antigens (10). The disease course seen in *B. rossi* infections bears a striking resemblance to that seen in falciparum malaria, leading one to hypothesize that a similar ‘cytokine storm’ may be an essential mechanism in the pathogenesis of this disease (11, 12). It is clear that *B. rossi* initiates a marked inflammatory response characterized by increased circulating markers such as C-reactive protein (CRP) and cytokines including monocyte-chemotactic protein-1 (MCP-1), interleukin (IL)-2, IL-6, IL-10, IL-18 and tumor necrosis factor alpha (TNFα) (13–15). Complications associated with this disease are likely the result of an unbalanced inflammatory cytokine response (13, 14, 16–18). Additional clinical and systemic indicators of inflammation, used to monitor affected dogs, which are associated with poor outcome in *B. rossi* infections include increased band neutrophil count, clinical collapse, presence of cerebral neurological signs and high parasitaemia (2, 14, 19).

Although we have some understanding of the inflammatory response triggered by *B. rossi*, all the existing research was performed in natural infections in dogs of various breeds, presented at variable disease stages and severity. In this prospective longitudinal experimental study, we aimed to investigate changes in markers of inflammation (cytokine concentrations, neutrophil count, monocyte count and CRP) and indicators of disease severity (including habitus, appetite, vital parameters, parasitaemia and hematocrit) over time in an experimental *B. rossi* infection of beagle dogs. We also aimed to investigate the influence infectious dose and treatment would have on disease progression. Finally, we wanted to identify if any significant correlations existed among the markers of inflammation and indicators of disease severity. We hypothesized, and found, that *B. rossi* infection initiates a pronounced, unbalanced inflammatory response and that infectious dose as well as treatment alter disease progression.

## Materials and methods

### Animals

This prospective longitudinal experimental study included six 6-month-old purpose bred sterilized male beagle dogs. The dogs were obtained from a commercial breeder (StudVet Beagles, RSA) and microchipped to allow for accurate identification. Vaccination and deworming programs were current. All dogs were clinically healthy and free of regional tick-borne diseases at the start of the experimental study, confirmed by hematology, serum biochemistry and polymerase chain reaction and reverse line blotting (PCR-RLB) for *Babesia, Ehrlichia, Theileria* and *Anaplasma*. One dog was splenectomized and used to raise a viable parasite inoculum from cryopreserved wild type *B. rossi*. The remaining five dogs were randomly assigned to one of two groups, namely the 3 dogs in the high dose (HD) and 2 dogs in the low dose (LD) groups, and experimentally infected with the corresponding *B. rossi* parasite inoculum dose. The mean of samples collected at two separate time points, 4 and 2 weeks prior to inoculation, from each of the 5 remaining experimental dogs formed the baseline data against which changes were compared overtime. Samples from the dog selected for the splenectomy were not included in the baseline data set. This experimental study was approved by the Animal Ethics Committee of the Faculty of Veterinary Science at the University of Pretoria (REC048-19).

### Preparation of the *Babesia rossi* inoculum and initiation of the infection

One randomly selected dog was splenectomized by a specialist surgeon and allowed 4 weeks recovery time after the surgery. The cryopreservate was created using blood from a dog naturally infected with *B. rossi* which was tested, and found to be negative for other blood-borne parasites using PCR-RLB and stored at −80°C. The splenectomized dog was then injected with 2 mL of thawed *B. rossi* cryopreservate intravenously, followed by a further 2 mL 24 hours later. Parasitemia was determined 12-hourly using a previously described technique, starting one day post-inoculation (19). Parasitemias were all determined on central venous blood. Once a parasitemia was detected and quantified, citrated whole blood was collected from the splenectomized dog. Using culture media (Culture Media RPMI 1640, Hepes, filtered water, sodium bicarbonate, sodium pyruvate and gentamycin), the blood sample was serially diluted to obtain the two inoculum doses, 10^8^ and 10^4^ parasitized red blood cells for the high and low doses respectively. The dogs in the HD and LD groups were then inoculated intravenously with the respective doses.

### Daily monitoring

All dogs were examined by a veterinarian once daily from the day of inoculation until the onset of clinical signs and thereafter as frequently as was deemed necessary to ensure adequate care until recovery. The experiment lasted a total of 8 days from point of inoculation to termination, with treatment being required from day 4. Habitus, appetite, temperature, heart rate, respiratory rate, mucous membrane color and blood pressure (using a non-invasive oscillometric technique – Vet-HDO^®^ Monitor) were determined daily, at the same time each morning. Blood pressure was measured on the tail of each dog, whilst lying in lateral recumbency. All dogs were thoroughly acclimated to this process prior to initiation of the experimental study to reduce stress associated increases in blood pressure during handling and sample collection.

### Hematology and biochemistry

Blood was collected atraumatically from the jugular vein into EDTA Vacutainer Brand Tubes (Beckton Dickinson Vacutainer Systems, UK) for a daily CBC (ADVIA 2120i, Siemens, Germany) and the EDTA plasma was then stored at −80°C. Blood samples were collected into serum Vacutainer brand tubes (Beckton Dickinson Vacutainer Systems, UK) every second day for CRP measurements. The CRP was analyzed using canine specific immunoturbidimetric CRP method^h^ (Gentian, Norway) run on the Cobas Integra 400 plus (Roche, Switzerland).

### Cytokine analysis

Once the experiment was concluded, the stored batched EDTA plasma samples were thawed at room temperature and used to determine granulocyte-macrophage colony-stimulating factor (GM-CSF), interferon gamma (INFγ), IL-2, IL-6, IL-7, IL-8, IL-15, IL-10, IL-18, TNF-a, INFγ-induced protein 10 (IP-10), keratinocyte chemoattractant (KC-like) and MCP-1. Concentrations were determined in duplicate by fluorescent-coded magnetic beads (MagPlex-C; MILLIPLEX. MAP Kit, Canine Cytokine Magnetic Bead Panel, 96-Well Plate Assay, CCYTO-90K, Millipore, Billerica, MA), based on the Luminex xMAP technology (Luminex 200, Luminex Corporation, Austin, TX). Two quality controls were included in the plate as internal quality controls. The assay was performed according to the manufacturer’s instructions. Cytokine concentrations were determined by comparing the optical density of the samples to the standard curves, produced from standards run on the same plate. The minimum detectable concentrations of the cytokines provided by the manufacturer were regarded as the detection limits in this study. Measurable values below the detection limit were assigned a value equal to the minimum detectable concentration for the respective cytokine and those with no measurable values were set as zero.

### Chemotherapeutic intervention

The infection was allowed to run its course until one of the following endpoints were identified: hematocrit <15%, collapsed habitus, nervous signs (such as seizure activity), clinical evidence of lung pathology with arterial blood gas evidence of acute respiratory distress syndrome (arterial partial pressure of oxygen [PaO2] <60mmHg), serum creatinine > 200mmol/L (normal <140 mmol/L) and hemoconcentration (PCV >55%). The infection ran its course for 4 days prior to intervention. The HD group was treated on day 4 in the morning (at 96 hours). Due to the unexpected death of one dog in the HD group, to avoid any further losses, the LD group was treated 12 hours later (at 108 hours) even though they had not reached the same degree of disease severity as the HD group. All dogs were drug cured with diminazene aceturate (3.5 mg/Kg subcutaneously) and provided with supportive treatment as needed. The remaining 5 dogs (including the splenectomized dog) recovered completely and were rehomed as pets.

### Statistical analysis

For the statistical analysis, variables that were shown to be non-normally distributed, were log-transformed; these were parasitemia, the leukocyte counts, CRP, GM-CSF, IFNγ, KC-like, all the interleukins, MCP-1 and TNFα. The other variables were not transformed. The means of the variables were then compared between the HD group and the LD group at each time point as well as between each time point and the mean baseline value within each group using linear mixed models, with animal identity as a random effect and the Bonferroni adjustment for multiple comparisons was applied. Pairwise correlations between variables were assessed using Spearman’s rank correlation. Significance was assessed at P<0.05. Statistical analysis was done using Stata 15 (StataCorp, College Station, TX, U.S.A.). Significant values in the text will be presented as the mean followed by the range and P value. Graphical presentation of some variables are included with error bars representing the standard deviation.

## Results

The demographic characteristics of the experimental group of dogs were as follows: All dogs were 6-month-old sterilized male beagle dogs. All 6 dogs tested negative for *Babesia, Ehrlichia, Anaplasma* and *Theileria* based on PCR-RLB done prior to the initiation of the experimental study. No significant difference was noted between the LD group and HD group for baseline data for any variable.

### Clinical variables

The HD group demonstrated changes in the clinical variables including habitus, appetite, temperature, heart rate and respiratory rate between 36 to 48 hours earlier than the LD group, indicating a more rapid onset of clinical disease in this group. Increases in diastolic (76 mmHg, range 74 – 77 vs baseline: 65 mmHg, range 63 – 67; *p* = 0.013), systolic (156 mmHg, range 145 – 161 vs baseline: 124 mmHg, range 122 – 125; *p* < 0.001) and mean arterial pressures (105 mmHg, range 101 – 110 vs baseline: 86 mmHg, range 74 – 87; *p* < 0.001) above baseline were noted in the HD group at 72 hours.

### Clinicopathological variables

The HD group had a detectable parasitemia 48 hours earlier than the LD group (Fig 1). There was a rapid increase in parasitemia thereafter, peaking at 46.76% (range 34.95 – 59.8) at 96 hours in the HD group and 5.76% (range 4.71 – 6.81) at 108 hours in the LD group. Parasitemia was strongly correlated to KC-like (*r* = 0.888, *p* < 0.001), IL-10 (*r* = 0.676, *p* = 0.009) and mature neutrophil count (*r* = −0.674, *p* < 0.001). Hematocrit (Hct) (Fig 2) declined significantly decline compared to baseline at 96 hours (*p* < 0.001) in the HD group and 120 hours (*p* = 0.003) in the LD group and both groups demonstrated a progressive decrease in Hct after treatment. Hemoglobinemia was visibly present from 72 hours in the HD group.

**Fig 1.**
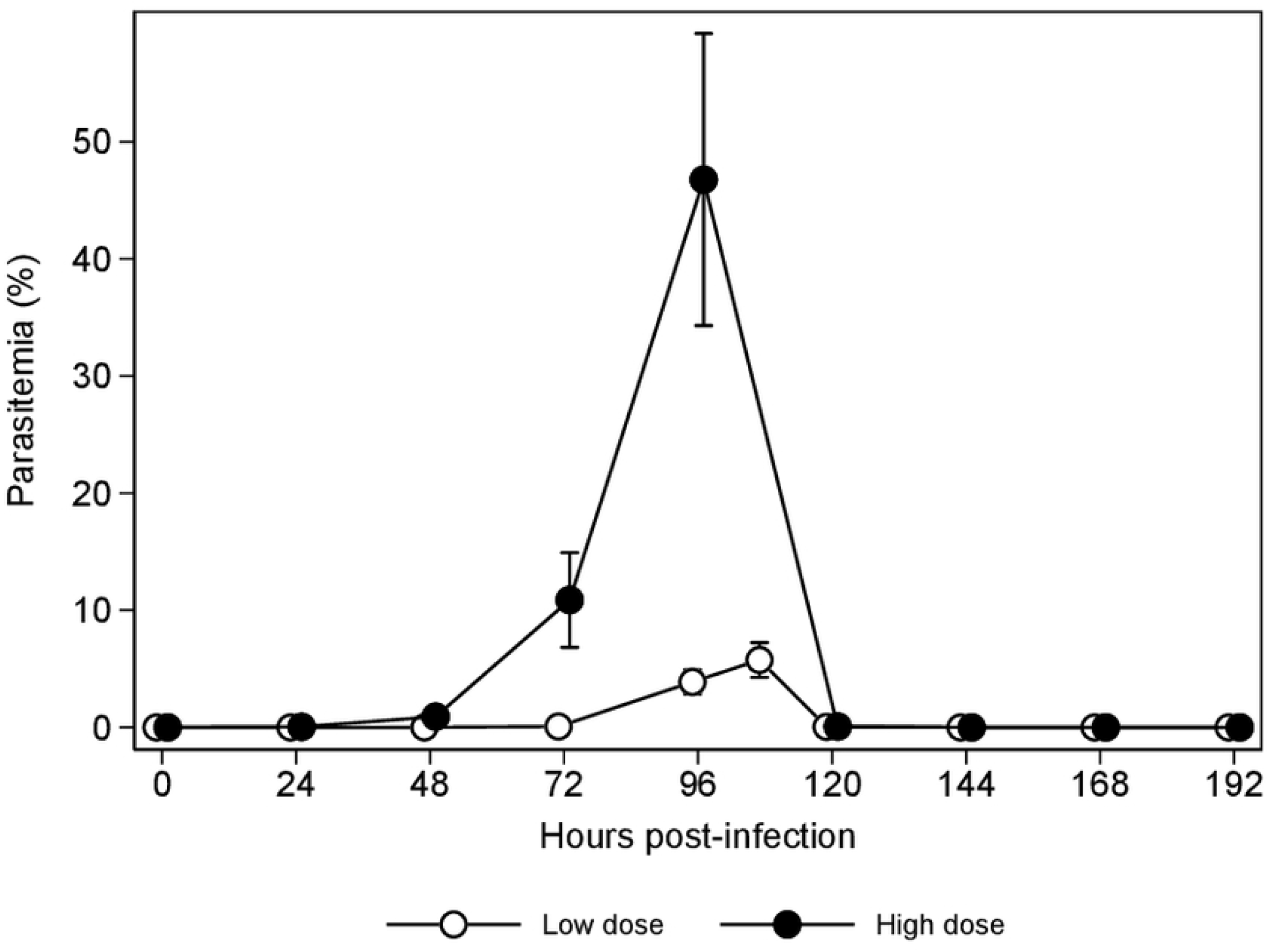
Parasitemia from inoculation of *B. rossi* until 4 days after treatment (Error bars represent there SD)

**Fig 2.**
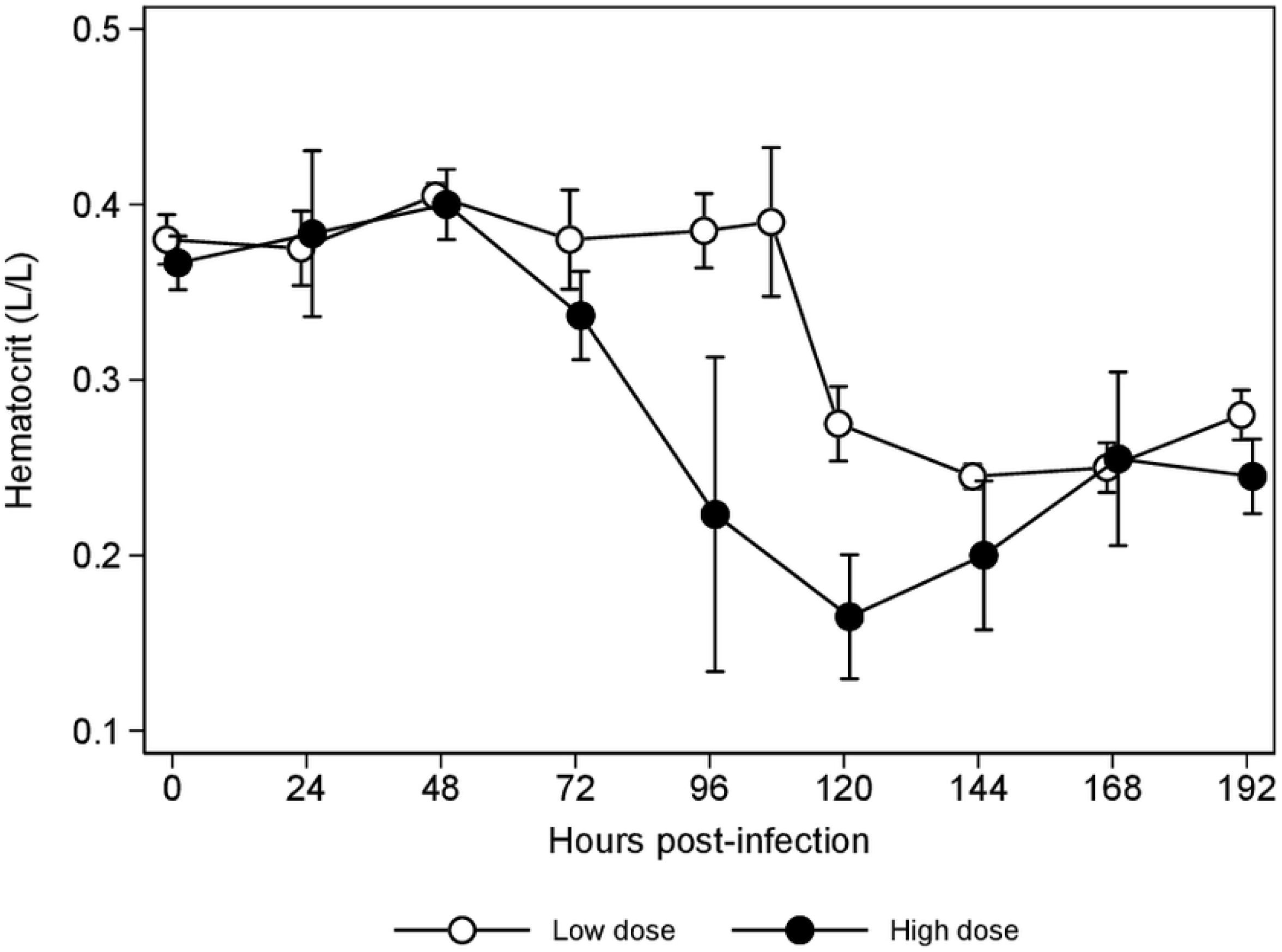
Hematocrit during *B. rossi* infection and after treatment (Error bars represent there SD)

For the HD group, increases in CRP concentrations (Fig 3) above baseline (14.33 mg/L, 10 – 21.5) peaked at 72 hours (150 mg/L, 135 – 163, *p* < 0.001) and remained significantly increased at 96 hours (125 mg/L, 92 – 160, *p* < 0.001), 120 hours (81.5 mg/L, 81 – 82, *p* < 0.001) and 144 hours (59 mg/L, 54 – 64,*p* < 0.001). The LD group showed a marked increase in CRP above baseline (25 mg/L, 10 – 40) at 108 hours (175 mg/L, 160 – 197, *p* < 0.001), declining thereafter but remaining significantly increased for the remainder of the study. The CRP concentrations peaked 36-hours earlier in the HD group. C-reactive protein was significantly correlated with temperature (*r* = 0.722, *p* = 0.003) and the correlation between CRP and parasitemia approached significance (*r* = 0.646, *p* = 0.056). A significant decrease in mature neutrophil (Fig 4) count was seen from 72 hours (1.79 × 10^9^/L, 1.36 – 2.44, *p* < 0.001) in HD group and 108 hours in the LD group (1.49 × 10^9^/L, 1.17 – 1.81, *p* < 0.001). The neutrophil nadirs for the HD and LD groups were seen at 96 (1.57 × 10^9^/L, 1.12 – 1.88, *p* < 0.001) and 108 hours (1.49 × 10^9^/L, 1.17 – 1.81, *p* < 0.001) respectively. In the HD group there was a marked increase in the mature neutrophil counts after treatment, exceeding laboratory reference intervals (3 – 11.5 × 10^9^/L) at 168 (17.64 × 10^9^/L, 12.21 – 23.07, *p* < 0.001) and 192 hours (27.35 × 10^9^/L, 23.75 – 30.94, *p* < 0.001). After treatment there was a gradual recovery of the mature neutrophil count in the LD group, returning to within laboratory reference intervals at 192 hours. A significant increase in band neutrophil counts (Fig 4) above baseline (0.14 × 10^9^/L, 0.13 – 0.17) was seen in the HD group at 120 hours (1.81 × 10^9^/L, 0.88 – 2.74, *p* < 0.001), 168 hours (2.77 × 10^9^/L, 2.28 – 3.25, *p* < 0.001) and 192 hours (6.45 × 10^9^/L, 4.32 – 8.57, *p* < 0.001). Band neutrophils counts exceeded the laboratory reference interval (0 – 0.5 × 10^9^/L) consistently after treatment in the HD group. At no point during the study did the band neutrophil count in the LD group increase significantly above baseline values or exceed the laboratory reference interval. A significant reduction in monocyte count compared to baseline (0.52 × 10^9^/L, 0.46 – 0.65) was seen in the HD group at 24 (0.28 × 10^9^/L, 0.2 – 0.32, *p* = 0.044) and 48 hours (0.27 × 10^9^/L, 0.2 – 0.34, *p* = 0.024). Following treatment, the monocyte counts were increased at 144 (1.86 × 10^9^/L, 1.27 – 2.45, *p* < 0.001), 168 (3.13 × 10^9^/L, 2.69 – 3.57,*p* < 0.001) and 192 hours (3.04 × 10^9^/L, 1.8 – 4.28,*p* < 0.001) in the HD group, exceeding the laboratory reference interval (0.15 – 1.35x 10^9^/L) from 144 hours onwards.

**Fig 3.**
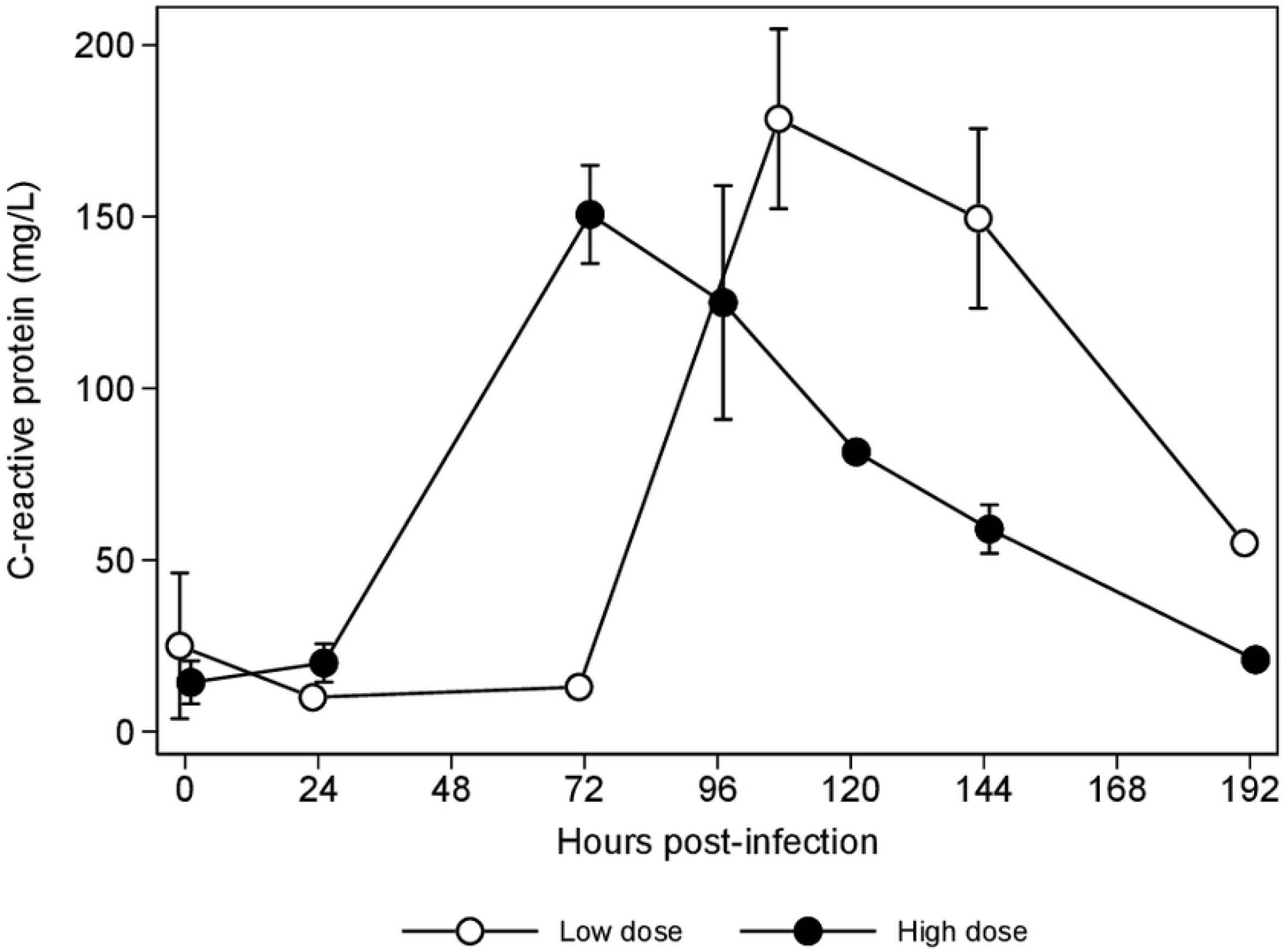
C-reactive protein concentrations during *B. rossi* infection and after treatment

**Fig 4.**
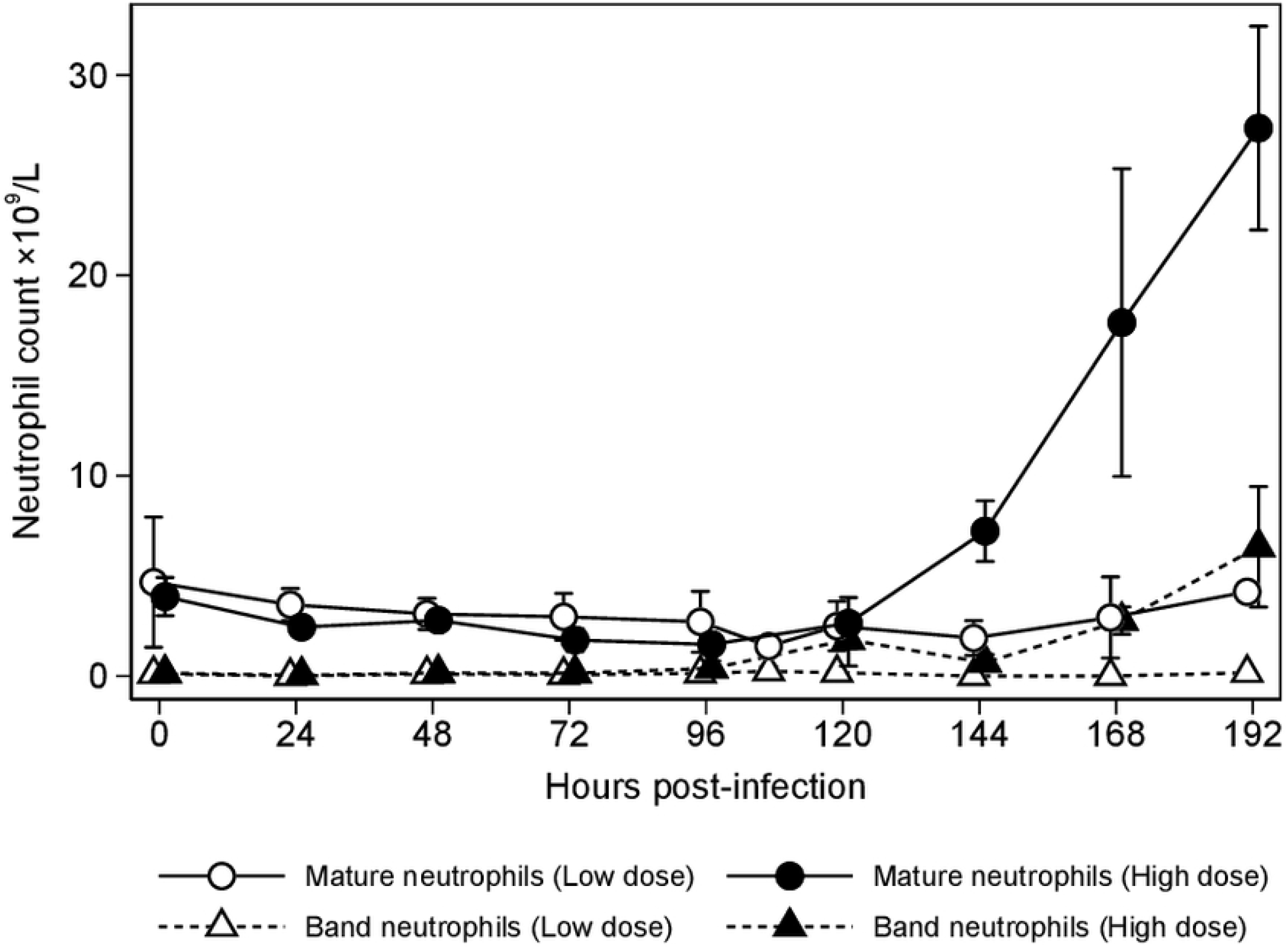
Mature and band neutrophil counts during *B. rossi* infection and after treatment

### Cytokine kinetics

Thirteen cytokines were evaluated, and the results were divided into 4 groups by pattern of change.

#### a. Cytokines that increased during infection and decreased after treatment

The cytokines which fall into this category included IFNγ and KC-like (Table 1). The HD group had a significant increase in IFNγ concentrations (Fig 5) above baseline (*p* = 0.002) and above the LD group (*p* < 0.001) at 48 hours. The LD group had peak concentrations 48-hours later, at 96 hours (*p* < 0.001). There was a progressive and significant increase in KC-like concentrations (Fig 6) in the HD group above baseline at 24 hours (*p* < 0.001), 48 hours (*p* < 0.001), 72 hours (*p* < 0.001) and 96 hours (*p* < 0.001) declining significantly at 144 hours (*p* = 0.004) and 192 hours (*p* < 0.001). The LD group only had significant increase in KC-like concentrations above baseline at 96 hours (*p* < 0.001). Strong correlations were identified between KC-like and parasitaemia (*r* = 0.888, *p* < 0.001) as well as KC-like and mature neutrophil count (*r* = −0.817, *p* < 0.001).

**Fig 5.**
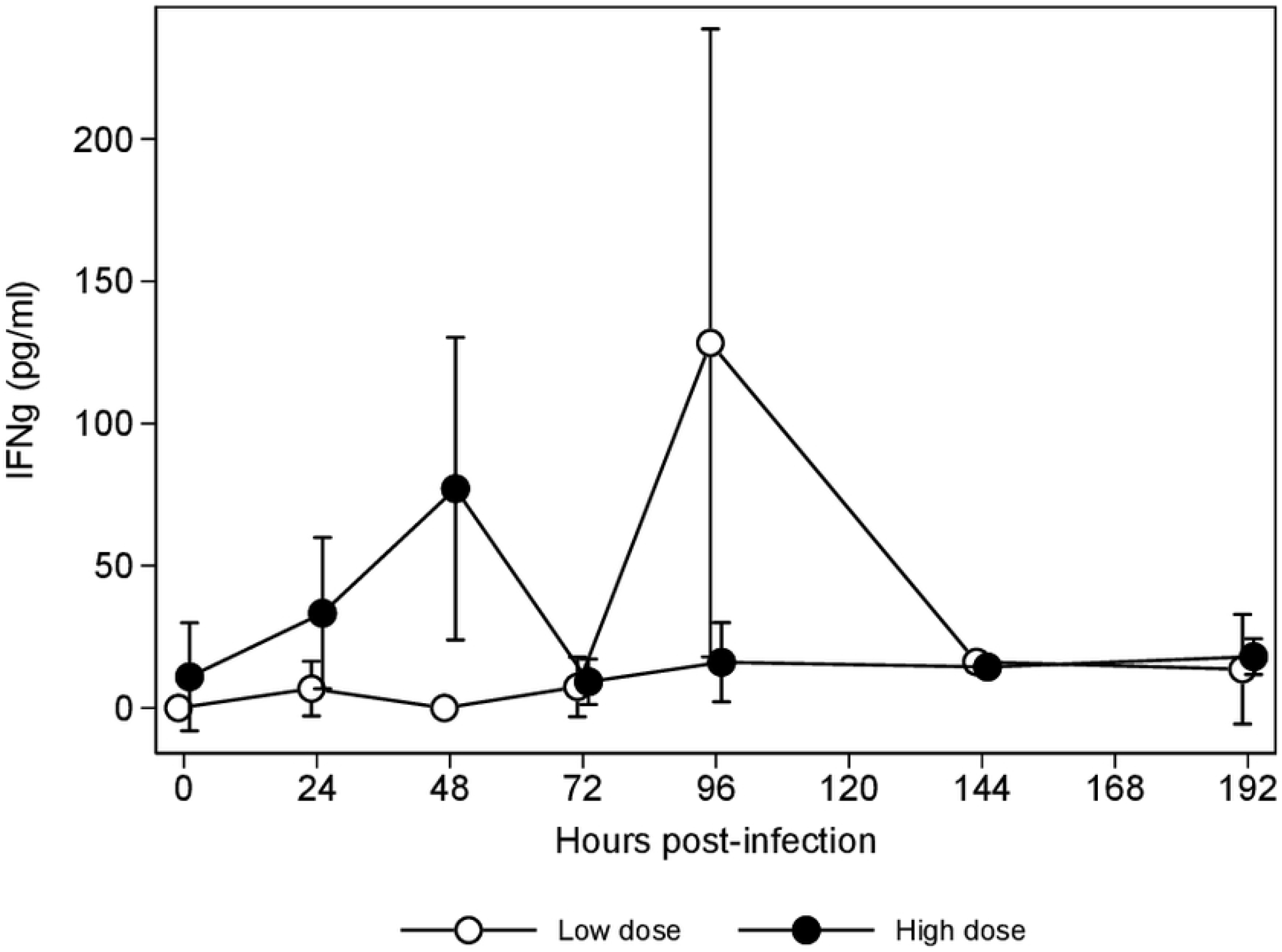
IFN γ concentrations during *B. rossi* infection and after treatment

**Fig 6.**
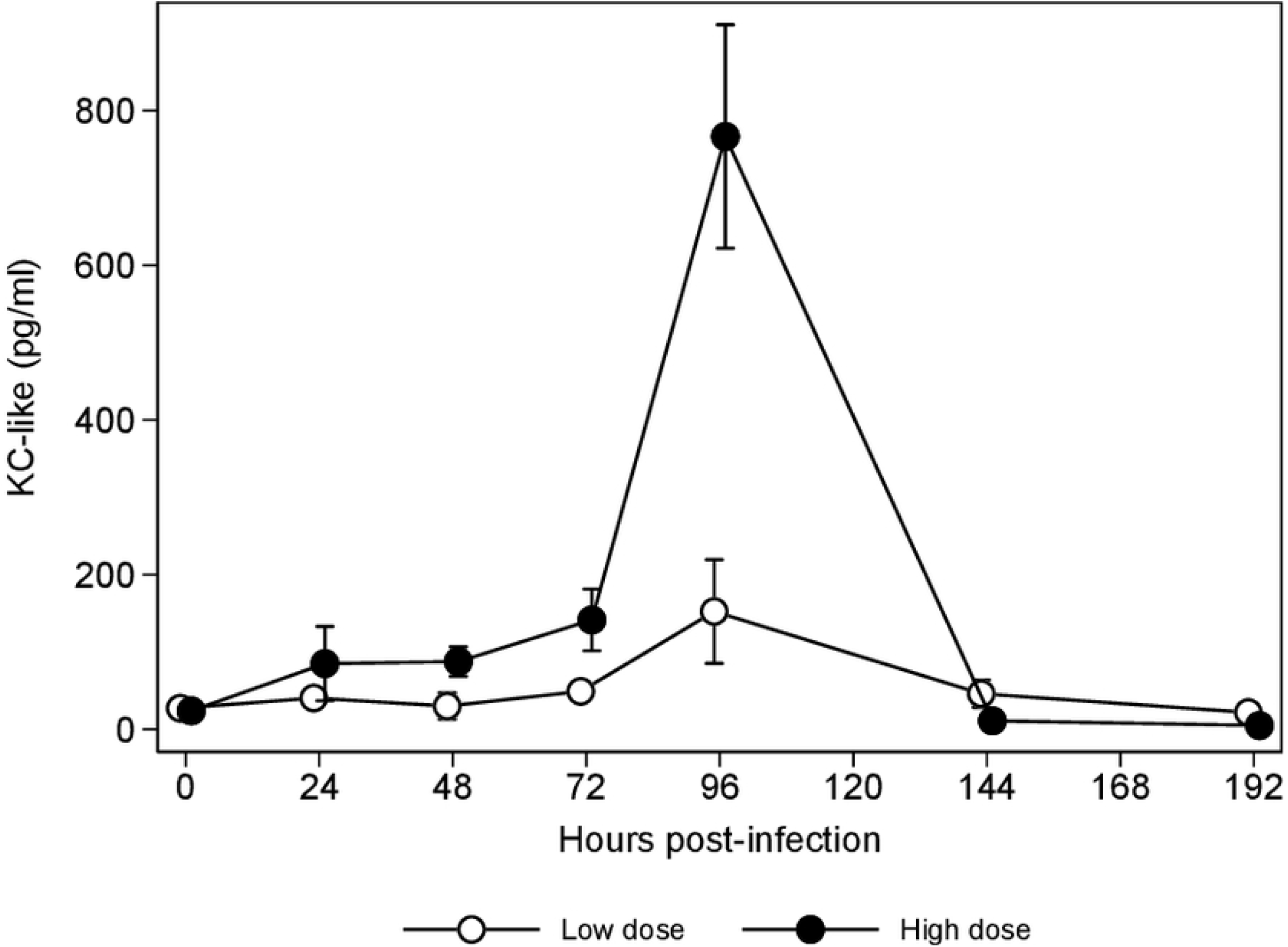
KC-like concentrations during *B. rossi* infection and after treatment

**Table 1:**
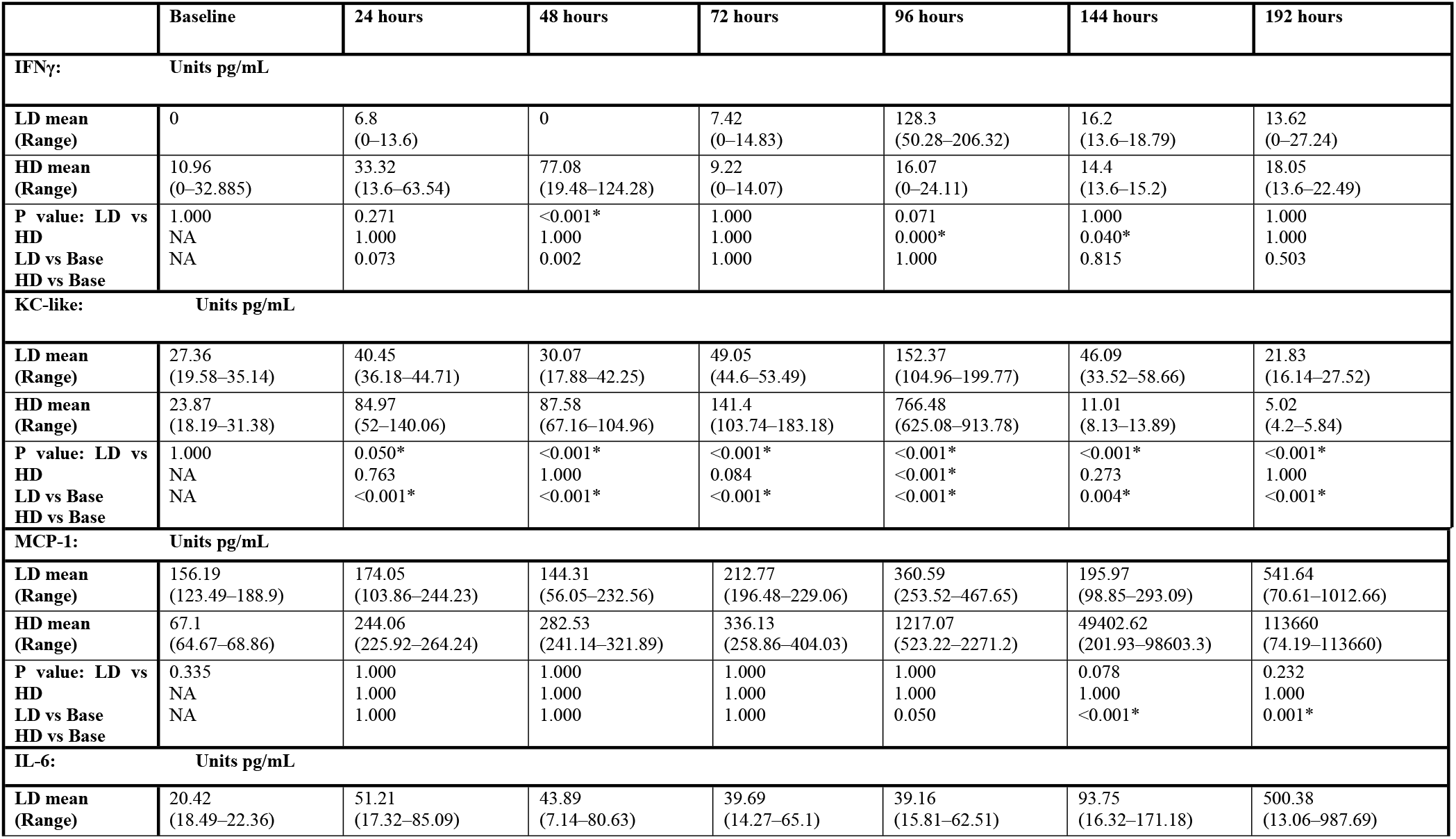
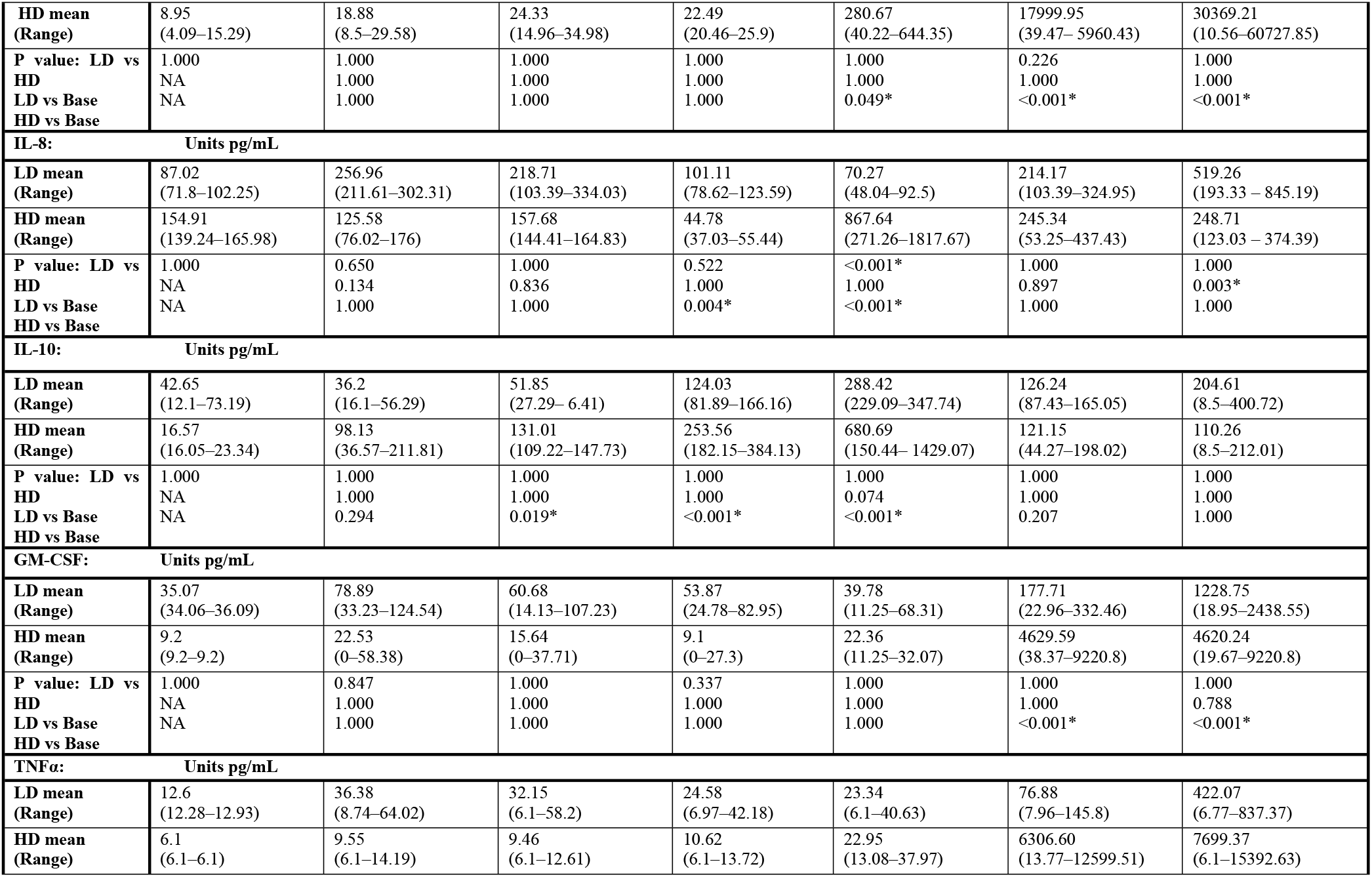
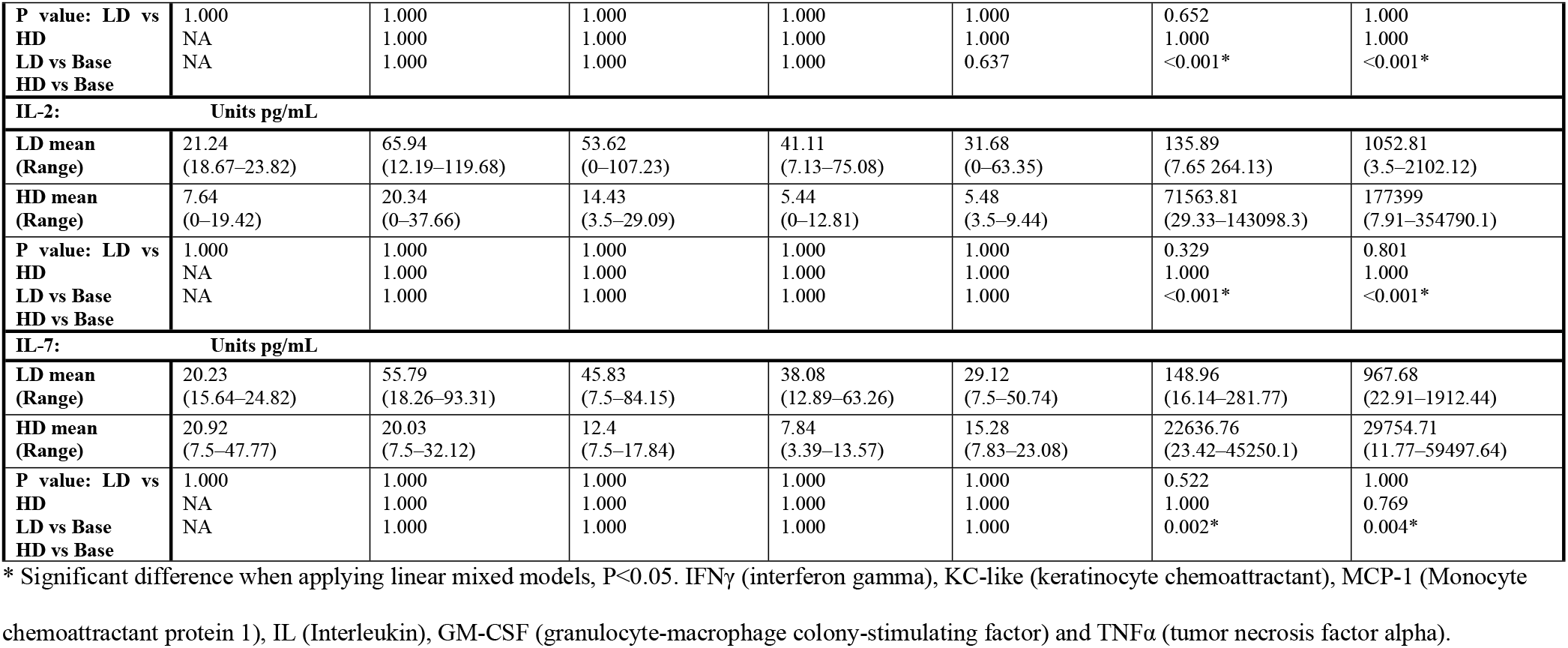
Cytokine concentrations during *B. rossi* infection and after treatment

#### b. Cytokines that increased during infection and remained high after treatment

This category included the following cytokines MCP-1, IL-6, IL-8 and IL-10 (Table 1). The chemokine MCP-1 concentrations (Fig 7) increased above baseline in the HD group from 24 hours onwards reaching significance after treatment at 144 hours (*p* < 0.001) and 192 hours (*p* = 0.001). The LD group did not have significant increases in MCP-1 concentrations throughout the study but one dog in this group did demonstrate increased concentrations after treatment. Interleukin-6 concentrations (Fig 8) were moderately increased above baseline at 96 hours (*p* = 0.049) in the HD group and progressively increased after treatment, at 144 hours (*p* < 0.001) and 192 hours (*p* < 0.001). Like changes seen in MCP-1, one dog in the LD group had increased IL-6 at 192 hours but this did not reach significance. Interleukin-6 and MCP-1 were strongly correlated (*r* = 0.792, *p* < 0.001). The HD group had significantly reduced IL-8 concentrations (Fig 9) compared to baseline at 72 hours (*p* = 0.004), with a marked increase at 96 hours (*p* < 0.001). The LD group only had significantly increased IL-8 concentrations at 192 hours (*p* = 0.003). Interleukin-10 concentrations (Fig 10) increased significantly above baseline at 24 hours (*p* = 0.019), 72 hours (*p* < 0.001) and 96 hours (*p* < 0.001) in the HD group. The LD group showed no significant increase in IL-10 but both dogs demonstrated a progressive increase in concentrations from 72 hours after inoculation until treatment, remaining increased in one dog after treatment. A strong positive correlation was identified between parasitaemia and IL-10 (*r* = 0.674, *p* = 0.009) as well as between IL-10 and MCP-1 (*r* = 0.828, *p* < 0.001). Similar kinetic profiles were seen between the HD and LD groups for MCP-1 and IL-10, varying in onset but not necessarily severity. IL-8 and IL-6 concentrations however appeared to follow different kinetic pathways between the HD and LD groups with the LD group demonstrating higher concentrations for the first 72 hours after inoculation. Interestingly the dog that died in the HD group had considerably higher concentrations of MCP-1 and IL-6 than any other dog at 96 hours.

**Fig 7.**
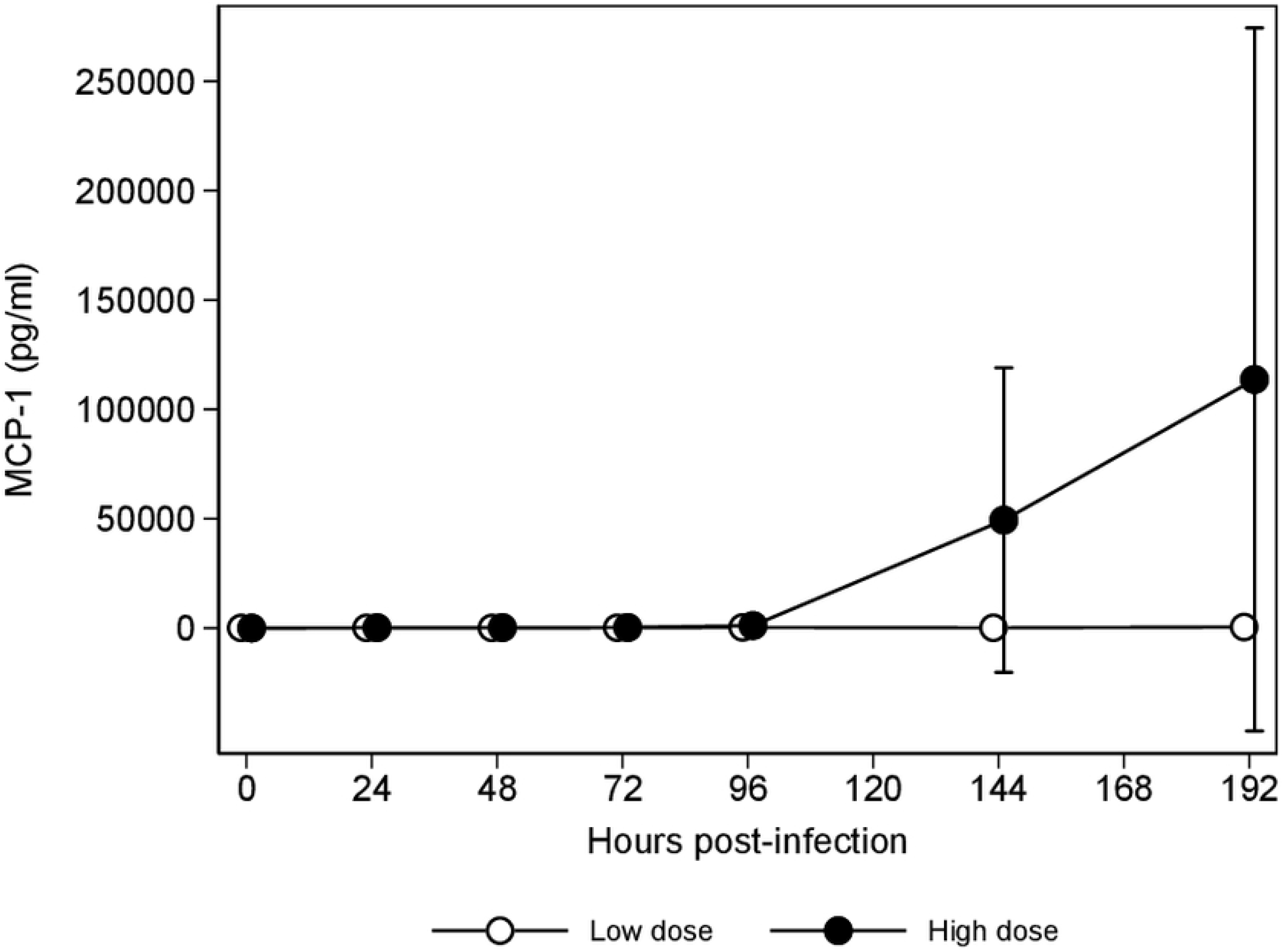
MCP-1 concentrations during *B. rossi* infection and after treatment

**Fig 8.**
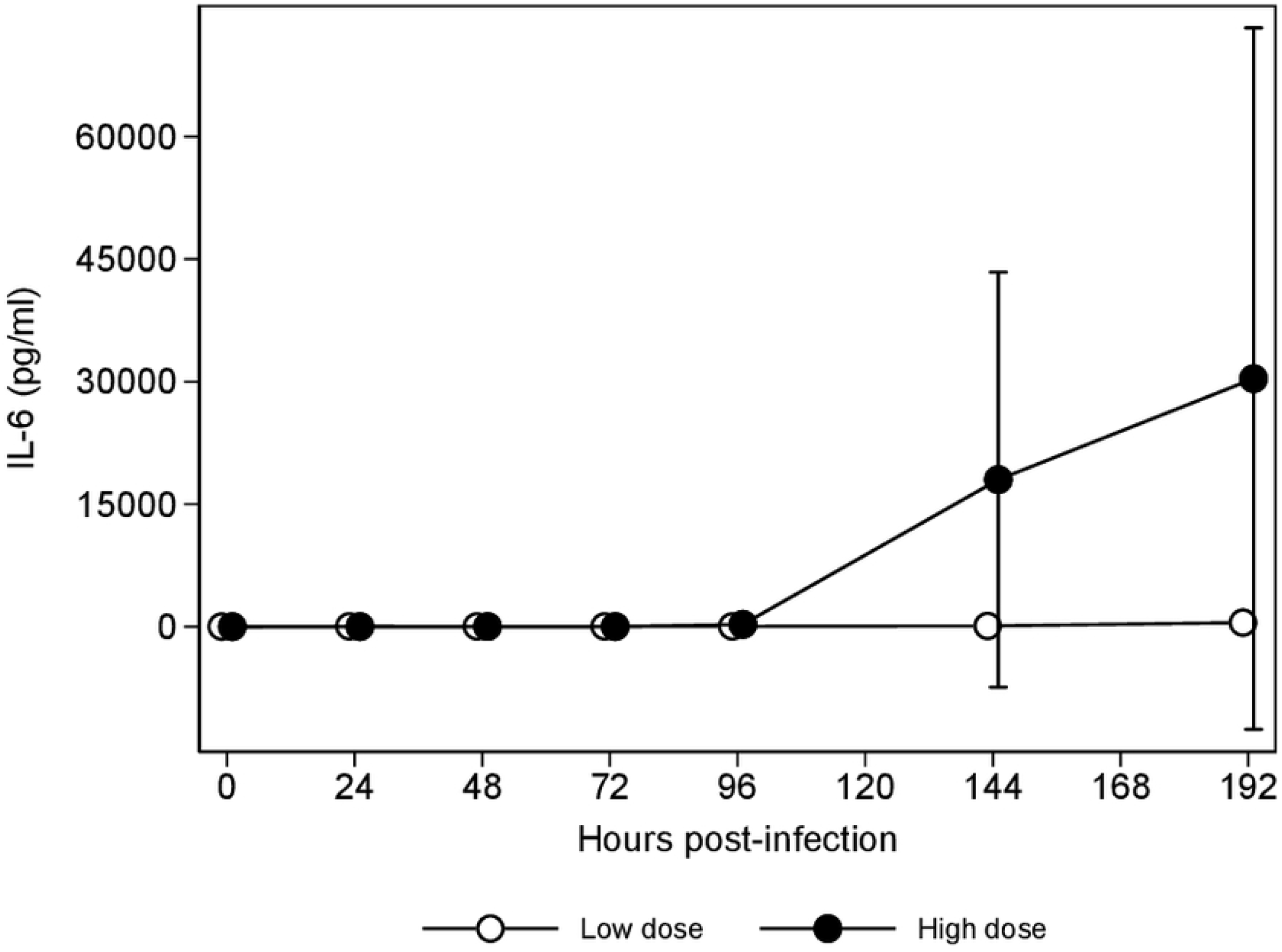
IL-6 concentrations during *B. rossi* infection and after treatment

**Fig 9.**
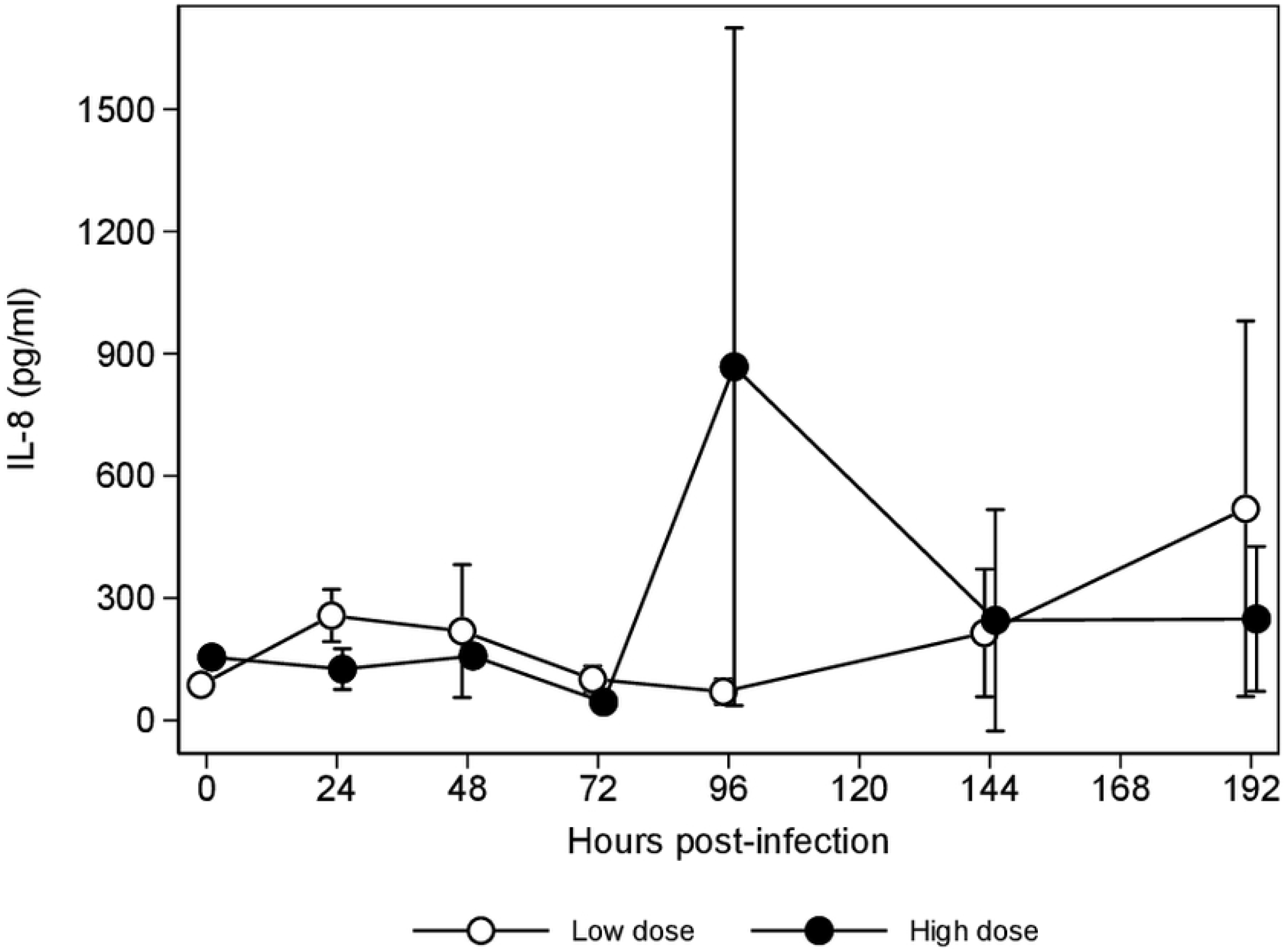
IL-8 concentrations during *B. rossi* infection and after treatment

**Fig 10.**
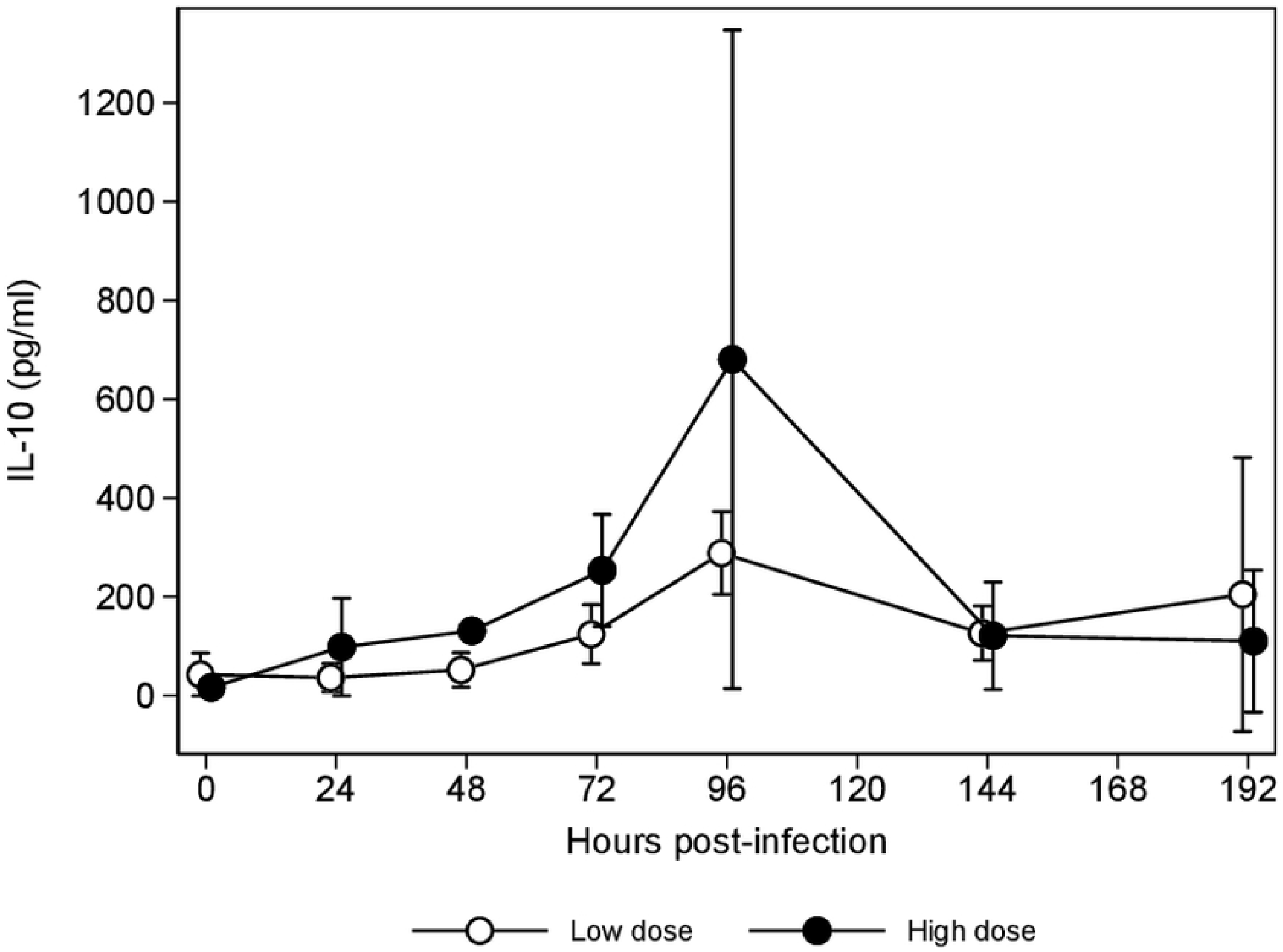
IL-10 concentrations during *B. rossi* infection and after treatment

#### c. Cytokines that increased after treatment

The next category of cytokines, GM-CSF (Fig 11), TNFα (Fig 12), IL-2 (Fig 13) and IL-7, had very similar patterns of change and were all markedly increased after treatment in the HD group, particularly in one dog (Table 1). Significant increases in these cytokines were seen after treatment, at 144 (GM-CSF *p* < 0.001; TNFα *p* < 0.001; IL-2 *p* < 0.001 and IL-7 *p* = 0.002) and 192 hours (GM-CSF *p* < 0.001; TNFα *p* < 0.001; IL-2 *p* < 0.001 and IL-7 *p* = 0.004). Although not statistically significant, one dog from the LD group showed a similar profile after treatment, with marked increases in all 4 cytokines although not to the same degree as seen in the HD group. The dogs with the highest parasitemia in each group demonstrated the greatest increases in cytokine concentrations after treatment. Tumour necrosis factor alpha demonstrated strong correlations with IL-6 (*r* = 0.925, *p* < 0.001), GM-CSF (*r* = 0.811, *p* < 0.001), IL-2 (*r* = 0.810, *p* < 0.001) as well as IL-7 (*r* = 0.810, *p* < 0.001). Interleukin 2 and IL-7 were also strong correlated (*r* = 0.872, *p* < 0.001).

**Fig 11.**
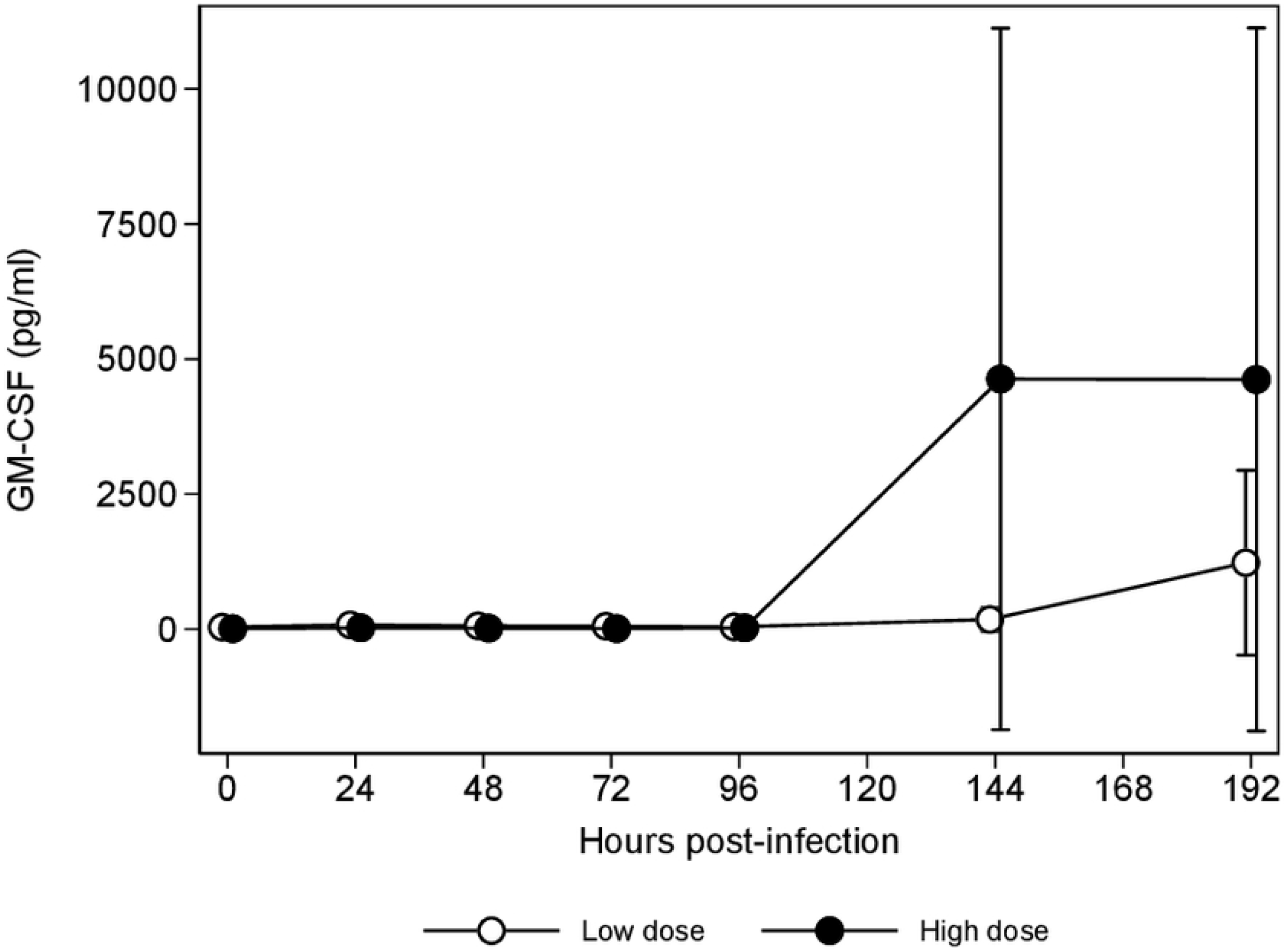
GM-CSF concentrations during *B. rossi* infection and after treatment

**Fig 12.**
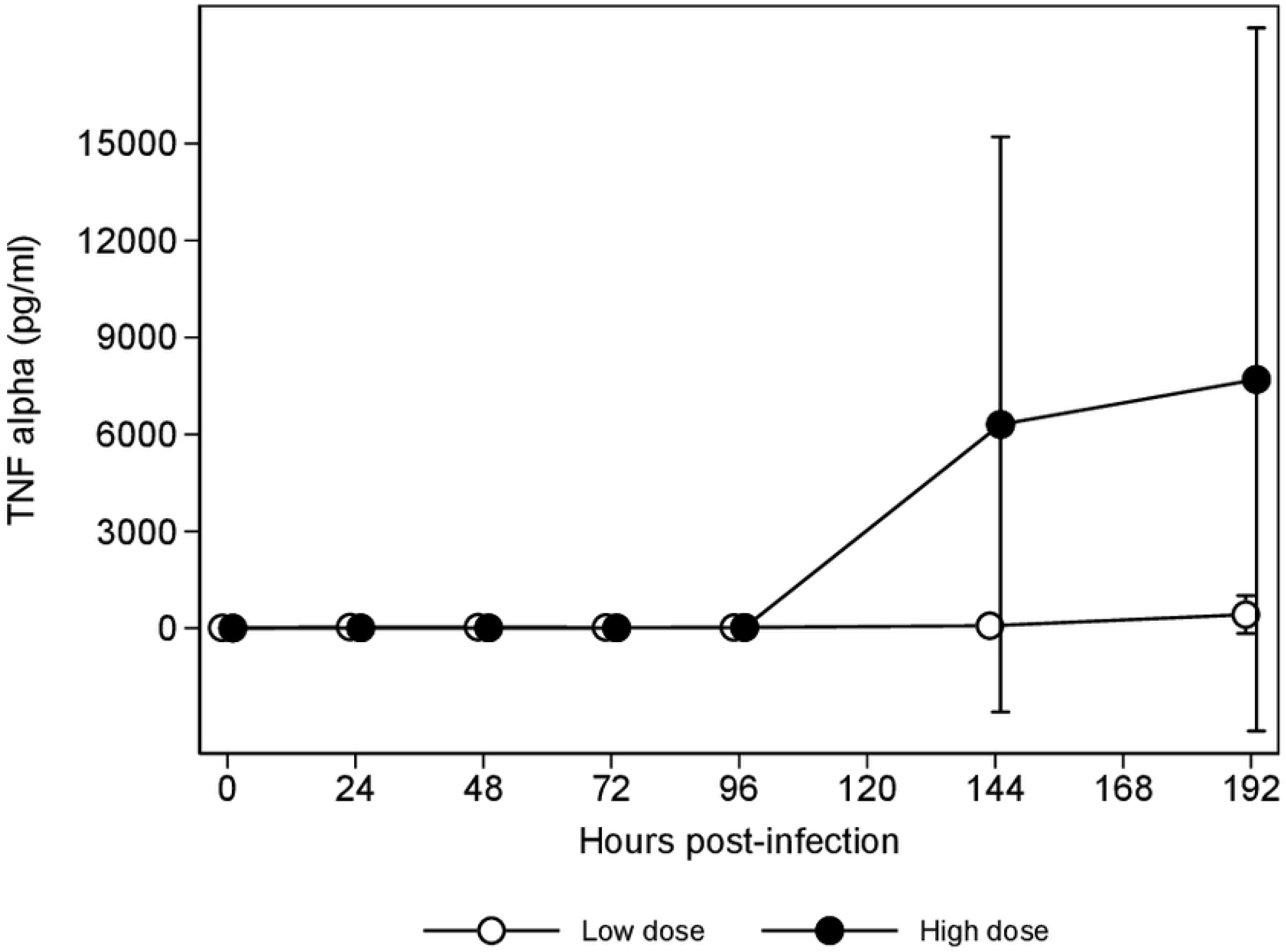
TNFα concentrations during *B. rossi* infection and after treatment

**Fig 13.**
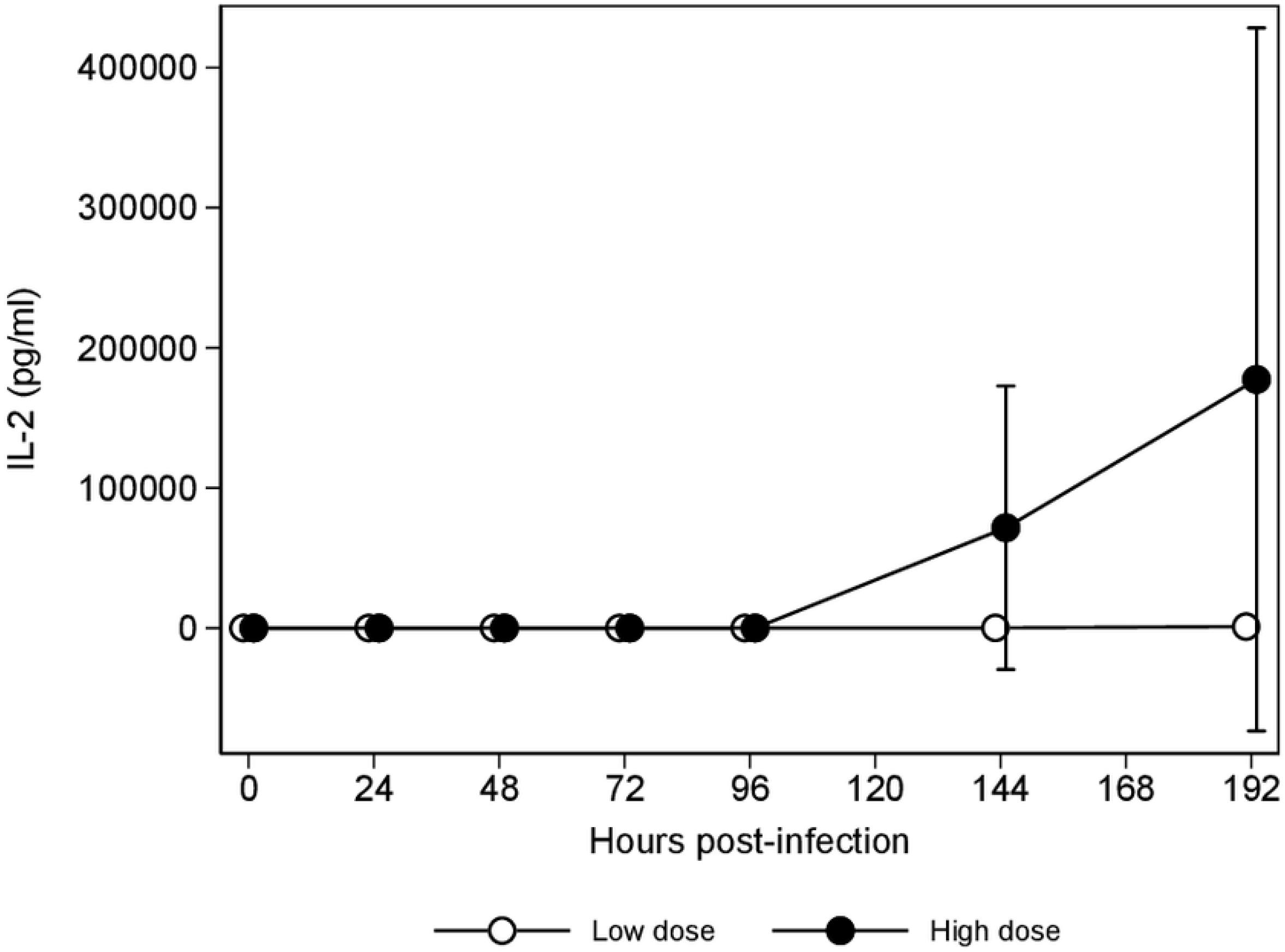
IL-2 concentrations during *B. rossi* infection and after treatment

#### d. Cytokines that showed no distinct pattern of change

The last three cytokines, IL-15, IL-18 and IP-10 showed minor changes in their concentrations during the course of the experiment. IL-15 (LD: 3043.38 pg/mL, 64 – 6022.76 vs HD: 23.36 pg/mL, 0 – 46.76, *p* < 0.001) and IL-18 (LD: 792.75 pg/mL, 18.86 – 1566.64 vs HD: 7.59 pg/mL, 0 – 15.17, *p* = 0.016) concentrations were significantly increased at 192 hours in the LD group. Interleukin 15 and IL-18 were strongly correlated (*r* = 0.981, *p* < 0.001). Finally, IP-10 showed mild increases in both groups during the study period.

## Discussion

Our study was the first to evaluate kinetics of markers of inflammation over the course of infection and recovery in a canine model experimentally infected with *Babesia rossi*. The results of this study illustrate the key role cytokines play in initiating and perpetuating inflammation in this disease and has also demonstrated that a pronounced inflammatory response continues and may even worsen despite clearance of the parasitemia. In addition to these findings the influence of inoculum dose was demonstrated, with a high infectious dose leading to an earlier onset of disease development and resulted in a more fulminant form of the disease. All the findings agreed with our original hypotheses, providing some insights into the pathogenesis of this hemolytic disease and even shedding additional light on the influence of treatment on the progression of the inflammatory response.

A progressive decline in both habitus and appetite was associated with higher infectious dose, rising parasitemia and worsening disease, improving with resolution of inflammation. As such, these clinical parameters can act as good indicators of disease onset and resolution. Although not statistically significant, there was a tendency for rectal temperature to increase with increasing parasitemia. The inoculum dose influenced the onset of changes in temperature, heart rate and respiratory rate in this study, with the HD group demonstrating increases 24 to 36 hours earlier than the LD group. Similar findings were seen in the experimental infection of dogs with *B. canis* (20). One dog in the HD group demonstrated a marked reduction in rectal temperature of 2.8°C at 96 hours, followed shortly thereafter by collapse and death. In a recent study of dogs naturally infected with *B. rossi*, rectal temperatures were significantly lower in dogs that presented collapsed with both collapse and hypothermia being positively associated with an increased risk of death (2). Rectal temperature may act as a proxy for the onset and progression of inflammation as well as the onset of hypoperfusion and shock. The blood pressure changes seen in our study differed from those seen during an experimental infection of dogs with *B. canis*, where the mean arterial blood pressure, measured using a non-invasive oscillometric blood pressure meter, declined progressively after inoculation (20). Mild hypotension was only identified in the one dog that died at 96 hours from the HD group. In previous studies hypotension worsened with disease severity (20, 21). The absence of hypotension in the remaining dogs in our study may be due to the short duration of the experiment and the timing of treatment. If the infection had been allowed to progress beyond our endpoints, we may have identified hypotension in more dogs.

High parasitemia is positively associated with increased risk of complications and death in *B. rossi* infections (14, 19). Venous parasitemia’s up to 30% have been recorded in natural *B. rossi* infections and this may be a contributor to the virulence of this *Babesia* species (19, 22). As with previous studies on *B. rossi* infection, we demonstrated a progressive parasitemia which required chemotherapeutic intervention. The infectious dose had a prominent impact on the progression of parasitemia with levels increasing at a significant rate and to very high levels in the HD group, up to 59%, within 4 days of inoculation. The LD group demonstrated a more gradual rise in parasitemia, more closely mirroring natural infection. The immune system may be overwhelmed and unable to mount an effective and timeous response to the *B. rossi* parasites when faced with high infectious doses or it is possible that these parasites actively suppress effective immune responses which may be more effectively suppressed at a higher parasitemia. The concept of an ineffective immune response may be supported by the positive correlation identified between parasitemia and IL-10, a prominent anti-inflammatory cytokine. Immune evasion by protozoa is a well-known phenomenon and has been demonstrated in *Plasmodium, Trypanosoma* and *Leishamania* among others (23). In *Leishamania* infections, the parasites promote an immunosuppressive cytokine profile with high levels of IL-10, allowing them unrestricted replication (23) and a similar interaction may take place in *B. rossi* infections. Immune dysregulation with concurrent hyperinflammation and immunosuppression is ubiquitous in human patients that are critically ill (24–26). It is also possible that, as in sepsis in humans, there is a state of hyperinflammation that is ineffective at clearing the infection but nevertheless does damage the host. The negative correlation between parasitemia and mature neutrophil count may also point to a deficient innate immune response to the *B. rossi* infection in these dogs. In addition to low neutrophil numbers a recent study on neutrophil function in *B. rossi* infections also identified an association between higher concentrations of neutrophil myeloperoxidase concentrations and poor prognosis suggesting possible diminished neutrophil burst function in the remaining neutrophils (27). It has also been shown that *B. rossi* results in significant lymphopenia as a result of a loss of CD3^+^, CD4^+^, CD8^+^ and CD21^+^ phenotypes (28). The degree of lymphocyte loss was also correlated to disease severity, and this may be responsible for a state of immune dysfunction despite hyperinflammation (28). Timing of treatment relative to parasitemia concentrations may have a marked influence on the degree of cytokine response after treatment, with cytokines such as TNFα, GM-CSF, IL-2 and IL-7 demonstrating marked increases after the parasites were damaged/killed by the treatment in the HD group with mild to moderate increases in the LD group. Higher parasitemia at the time of treatment may result in a more severe, unregulated pro-inflammatory response after treatment. Damage to the parasites releases soluble parasite antigens into circulation which could be efficient in stimulating this profound immune response. Although this response may increase the rate of *B. rossi* parasite clearance, it could be redundant and lead to unnecessary widespread ‘innocent bystander’ injury to host tissues.

Anemia is common in dogs infected with *B. rossi* and in a recent study up to 84% of dogs had hematocrits below the laboratory reference interval at presentation (2). Although anemia is not a reliable predictor of death, severe anemia does require treatment to avoid the systemic complications of hypoxia and even death (2). The HD group demonstrated a significant anemia at 96 hours after inoculation which worsened after treatment requiring multiple blood transfusions. The severity of the anemia after treatment in this group was likely underestimated as a result of this intervention. Although the decline in Hct was significant in the LD group after treatment, it only resulted in mild anemia in these dogs and blood transfusions were not necessary. The absence of anemia during active infection in the LD group was probably the result of insufficient time permitted for disease progression to reach the same severity as that seen in the HD group. A marked decline in hematocrit occurred after treatment in both groups similar to previous studies (29).

Mature neutropenia was identified as early as 24 hours post-inoculation in the HD group, but only reached statistical significance from 72 hours. The mature neutropenia persisted until treatment, thereafter, counts increased significantly above baseline and laboratory references intervals. A mature neutropenia was seen in the LD group, 36-hours after the HD group, but neutrophilia did not develop after treatment. Previous studies evaluating hematological changes in natural *B. rossi* infections found that a large percentage of dogs present with a neutropenia (27, 30). A progressive band neutrophilia was seen in the HD group after treatment and band neutrophilia in *B. rossi* infections has been associated with lower hematocrits and blood transfusions, consistent with findings in our study (30). A band neutrophil count of > 0.5 × 10^9^/L at presentation carries an odds ratio for death of 5.9 (2). Interestingly the only dog with a band neutrophil count above this level prior to treatment in our study was the dog that died. In one study there was a higher proportion of dogs with a neutrophilia in the group that received blood transfusions (30). The blood transfusions received by the HD group may have contributed to the left shift neutrophilia but hemolysis and subsequent increases in cytokine release and systemic inflammation are likely the main role players. There was a strong negative correlation between mature neutrophil count and KC-like, a cytokine with a major role in neutrophil migration and activation (31). Interleukin-8, another important cytokine in the migration and activation of neutrophils, had a peak concentration at 96 hours in the HD group, coinciding with the mature neutrophil nadir (32). The migration of neutrophils, under the influence of cytokine cues, out of circulation to various sites of inflammation may contribute to the circulating neutropenia seen in *B. rossi* infections. The cytokine, GM-CSF, stimulates the initiation of granulopoiesis in the bone marrow, and this cytokine increased after treatment, particularly in the HD group, coinciding with increases in neutrophil and monocyte counts (33). The HD group developed a mild monocytosis after treatment, similar to natural *B. rossi* infections (30). It should be noted that *B. rossi* infected dogs demonstrate monocyte/macrophage accumulation in the pulmonary interstitium and spleen (34, 35). A large increase in MCP-1 after treatment in the HD group indicates increased demand for monocyte/macrophage activity during this period and GM-CSF may have provided the bone marrow stimulation to increase production of monocytes after treatment (33, 36).

Acute phase proteins are regularly used in the detection and monitoring of systemic inflammation. C-reactive protein is an acute phase protein that is consistently elevated in canine babesiosis despite levels not correlating with outcome (15, 20, 37). In one study of CRP in natural *B. canis* infection, CRP had its peak concentration at presentation and declined progressively following treatment (37). In a *B. canis* experimental infection, CRP increased before the presence of a detectable parasitemia and the onset of the increase was inoculum dose dependent with the highest dose resulting in increased concentrations first (20). The inoculum dose in our study influenced the onset of increases in CRP concentration with the HD group showing a significant increases 36 hours earlier than the LD group. Low levels of parasitemia were detectable prior to significant increases in CRP concentrations in both groups unlike findings in experimental *B. canis* infection (20). This may however not have been the case had lower infectious doses been used. Treatment resulted in a progressive decline in CRP concentrations in both groups. Although CRP increased with parasitemia the correlation wasn’t statistically significant in this small cohort, but rectal temperature and CRP were positively correlated. As seen in the *B. canis* experimental study, CRP concentrations in our study reached a ceiling and remained relatively stable despite progressive parasitemia (20). C-reactive protein levels remained high even after parasitemia was undetectable and this delay was most likely due to the half-life (which is approximately 161 hours in dogs, with significant inter-individual variation) rather than continued production (38).

Cytokines are a group of proteins secreted by cells of the immune system which act as key signalling molecules in any inflammatory response. A number of cytokine changes have been identified in *B. rossi* infections, but these have only been evaluated in dogs at presentation, providing a single snap shot in time of a complicated and dynamic disease (13, 14). Cytokines shown to increase during *B. rossi* infections include IL-6, IL-10, MCP-1 and TNFα, and their concentrations tended to be higher in dogs with complicated disease (13, 14). Only IL-6 and IL-10 concentrations were significantly higher in dogs that died compared to survivors (13). Decreased concentrations of IL-8 were consistently identified in natural *B. rossi* infections, in contrast to *B. canis* infections (13, 14, 16).

Interferon gamma and KC-like increased with the start of infection and declined after treatment. These cytokines seem to be released by the host in an attempt to control the parasite biomass similar to the IFNγ response seen in falciparum malaria (39). In malaria, IFNγ is considered an important mediator in the protective innate immune response during the blood stage and initial parasite replication (40). It is possible that IFNγ is important in the initial immune response to *B. rossi* infection, suppressing early replication of the parasite as this cytokine increased early in the course of the experimental infection, coinciding with the initial increase in parasitemia in both groups. The concentrations, however, declined acutely once the parasitemia exceeded 5% in HD group. The sudden decline in IFNγ concentrations in the HD group coincided with a marked increase in parasitemia. The high levels of parasitemia seen in the HD group may have induced a state of immune exhaustion (41). High concentrations of IL-10 may also have contributed to the sudden decline in IFNγ, thereby suppressing its secretion and contributing to the resultant unregulated parasite replication (42–44). Suppression of IFNγ secretion may be one mechanism employed by the parasite allowing unrestrained increase in parasite biomass and it is possible that a similar decline may have been seen in the LD group if infection had been allowed to evolve. Keratinocyte chemoattractant-like increases in *B. gibsoni* and *B. canis* infections, and high concentrations were able to differentiate complicated from uncomplicated *B. canis* cases (16). In our study KC-like increased progressively during infection and declined following treatment, correlating strongly to parasitemia. In addition to promoting neutrophil migration, KC-like may contribute to increased risk of complications because of enhanced neutrophil release of reactive oxygen species and neutrophil extracellular traps, important mechanisms by which host tissue may be damaged (45).

Cytokine increases after treatment could reflect a role in the ‘run-away’ inflammation that persists even after the initial trigger has been removed. Two pro-inflammatory cytokines, MCP-1 and IL-6 had a similar pattern of change and showed a strong positive correlation with one another. Both cytokines displayed progressive increases from the point of inoculation increasing markedly after treatment in the HD group. No significant increases were noted in the LD group throughout the study period, but concentrations of MCP-1 were trending upwards 24 hours prior to treatment. In previous studies on natural *B. rossi* infections MCP-1 and IL-6 were increased in infected dogs at presentation and higher concentrations were associated with increased risk of mortality (13, 14). The concentrations of MCP-1 and IL-6 in the dog that died were considerably higher than any other dog in our study just prior to treatment indicative of a negative prognosis. Monocyte chemoattractant protein-1 recruits and activates monocytes/macrophages and would be a vital host mechanism in the immune response to babesia parasites by amplifying inflammatory signals and enhancing phagocytosis of parasites and damaged erythrocytes (36). Persistently high levels however could contribute to an unregulated inflammatory response and increased risk of complications such as acute lung injury seen in some dogs that die as a result of *B. rossi* infection (34). A potent stimulator of IL-6 production is TNFα, and a strong positive correlation was detected between IL-6 and TNFα in this study. The role of IL-6 in septic conditions is poorly understood, although it does play a role in many pro-inflammatory activities such as stimulating the production of acute phase proteins like CRP from hepatocytes (well known to be raised in *B. rossi* infection), activation of lymphocytes and acting as a pyrogen (15, 46). Interleukin-6 is also thought to link inflammation with thrombosis in sepsis (46, 47). Widespread formation of microthrombi is a well-defined pathology in canine babesiosis, particularly in *B. rossi* infections, and increases in IL-6 may be an important trigger for this, contributing to increased risk of complications such as cerebral babesiosis, myocardial dysfunction and death (48–50). Interleukin-6 is also shown to play an important role in the acute endocrine response to infection which is well described and so typical of this disease (51–53).

In previous studies on the cytokine changes in *B. rossi* infections, IL-8 was decreased at presentation when compared to healthy control dogs (13, 14). In contrast to these findings, IL-8 increases in *B. canis* infections and even showed a progressive rise for at least 7 days after treatment (16). Our study demonstrated decreased IL-8 concentrations during the early stages of infection in the HD group followed by a considerable increase shortly before treatment, when parasitemia was very high. Interleukin 8 concentrations remained high after treatment in this group. A mild transient increase in IL-8 concentrations was noted in the LD group 24-hours after infection, followed by a progressive decline in concentrations until treatment. Once treated, IL-8 concentrations increased progressively until 192 hours in this group. The decline in IL-8 production in *B. rossi* infections is poorly understood but because this cytokine is not constitutively produced and requires inflammatory stimulus, it is possible that in the early stages of infection, before parasitemia and hemolysis are severe, there is insufficient stimulus (32). Suppression of IL-8 and the cytokines that stimulate its production (TNFα and IL-1) during the initial phase of infection may be, in part, due to high concentrations of IL-10 (32, 54). The final cytokine in this group, IL-10, is a prominent anti-inflammatory cytokine. Concentrations of IL-10 progressively increased over the course of the experimental infection and decreased gradually after treatment. High IL-10 concentrations have been noted in natural *B. rossi* infections (13, 14). Interleukin-10 is essential in the modulation of the inflammatory response and plays a key role in preventing excessive inflammation as well as promoting the resolution of inflammation once the inciting pathogen has been eliminated (54). Although the anti-inflammatory effects of IL-10 are critical, excessive or inappropriately timed production of IL-10 may prevent an effective immune response to an organism, allowing persistence or even unregulated replication in the host (54). This has been seen in *Leishmania spp*. and *Plasmodium spp*. infections in which high IL-10 concentrations can lead to fulminant fatal infections or chronic persistent infections (54). Human septic patients with continuous over production of IL-10 and high IL-10:TNFα ratio develop marked immunosuppression and have a higher risk of mortality (55). A strong positive correlation was seen between IL-10 and parasitemia supporting the notion that IL-10 may have a permissive effect on the replication of *B. rossi* parasites.

The cytokines GM-CSF, TNFα, IL-2 and IL-7 only increased significantly after treatment. Increased levels of GM-CSF have been identified in *B. rossi* infections, particularly in dogs presented earlier in the course of disease (13). The rise in GM-CSF concentrations after treatment noted in our study may act as a double-edged sword, on the one side replenishment of neutrophil and monocyte counts is essential but on the other, excess production, adhesion and activation of granulocytes and macrophages after the parasites are eliminated may contribute to widespread tissue damage (33). The initiator pro-inflammatory cytokine TNFα is one of the most studied cytokines in human medicine and is an important mediator in the protection against microbial infections (56). It can however lead to pathology in cases where there is disproportionate and dysregulated immune response to an infection by the host (56). It is also a potent stimulator of the production of other pro-inflammatory cytokines such as IL-1β, IL-6 and IL-8, serving as co-ordinator in the inflammatory response (46). Increased concentrations were found in natural *B. rossi* infections and higher levels were associated with increased risk of complicated disease and death (14). Concentrations of TNFα were only increased after treatment in this study and reached very high levels in one dog in the HD group. There was also a strong positive correlation with IL-6, GM-CSF, IL-2 and IL-7. The excessive production of TNFα following treatment may be indicative of a dysregulated immune response. Both IL-2 and IL-7 act on lymphocytes and were positively correlated with each other in this study (57, 58). Interleukin-2 did not increase when *B. rossi* infected dogs were compared to healthy control dogs in one study but higher concentrations were noted in infected dogs presented within 48 hours of clinical illness (13, 14). Increases in IL-7 has not been identified in previous studies of *B. rossi* infections (14). Both cytokines only displayed significant increases after treatment in the current study. Previously, significant reduction in T-helper lymphocytes and cytotoxic T-lymphocytes were identified in complicated *B. rossi* infections and decreased concentrations of cytotoxic T-lymphocytes was also noted in uncomplicated cases at presentation (28). In that study it was hypothesised that a state of functional immune suppression may be present, and this is supported by the apparent deficiency of cytokines involved in lymphocyte proliferation and activation identified in our study prior to treatment (28). Treatment and subsequent lysis of the parasites may have interrupted the immunosuppressive state and the release of soluble antigens was able to stimulate the adaptive immune response triggering production of these cytokines.

Three cytokines showed no distinct pattern of change or only demonstrated changes in one dog. A strong positive correlation was identified between IL-15 and IL-18 in this study. Both these cytokines only increased significantly in one dog in the LD group after treatment. A trend in the increase of IL-15 concentrations early on in disease course of *B. rossi* infection was identified in one study (13). In the current study IP-10 concentrations were mildly increased in both groups.

The main limitation of this study was the small sample size. Six dogs we were used in the study, with only 5 dogs being inoculated. Every attempt was made to exclude any confounding or influencing factors. All dogs were the same age, sex and breed with identical vaccination and deworming protocols. They were housed in the same isolation housing and outdoor facilities. Diet, training, experimental procedures, sample collection and human interaction was consistent between all dogs. Due to the small sample size, it is possible that significant differences between the groups may have been missed (type 2 error).

Our study has found that infectious dose influenced the onset and dynamics of the inflammatory response. Most variables shared similar kinetic patterns between groups, differing with respect to the timing of the onset of disease only. If the infection in the LD group had been allowed to progress, it is likely these variables would have reached similar degrees of change to those seen HD group. There were however exceptions, where kinetic patterns differed during infection between the two groups such as those seen in IL-8. Concentrations of CRP, KC-like, IL-15, IL-18 and IP-10 in the LD group exceeded those of the HD group after treatment. These findings suggest that not only will infectious dose influence the onset of inflammation, but it may influence the kinetics and nature of the inflammatory response to *B. rossi* infection. Moreover, the level of parasitemia may be a contributor to the development of complications after treatment. We also highlighted the influence chemotherapeutic damage to the parasites had on the progression of the inflammatory response. We found that not only was treatment unsuccessful in curbing the inflammatory response, but it may augment it by triggering the production of several pro-inflammatory cytokines and proliferation of inflammatory cells. Progression of the inflammatory response after treatment would be redundant and may even lead to unnecessary host tissue damage. It is clear from the findings of our study that *B. rossi* infection and treatment triggers a classical ‘cytokine storm’ in which the host’s response is characterized by severe inflammation and tissue damage beyond that induced by the parasite itself (9, 59).

## Acknowledgements

**Melvyn Quan**

Department of Veterinary Tropical Diseases, Faculty of Veterinary Science, University of Pretoria, Pretoria, South Africa

Contributions: Assistance in the running of the cytokine analysis.

## Notes

### Competing Interest Statement

The authors have declared no competing interest.

